# BORCS6 is involved in the enlargement of lung lamellar bodies in *Lrrk2* knockout mice

**DOI:** 10.1101/2021.03.05.434068

**Authors:** Miho Araki, Sho Takatori, Genta Ito, Taisuke Tomita

**Affiliations:** Laboratory of Neuropathology and Neuroscience, Graduate School of Pharmaceutical Sciences, The University of Tokyo, Tokyo 113-0033, Japan; Social Cooperation Program of Brain and Neurological Disorders, Graduate School of Pharmaceutical Sciences, The University of Tokyo, Tokyo 113-0033, Japan

**Keywords:** BLOC-1-related complex subunit 6, leucine-rich repeat kinase 2, lung lamellar body

## Abstract

Leucine-rich repeat kinase 2 (LRRK2) has been implicated in the pathogenesis of Parkinson disease. It has been shown that *Lrrk2* knockout (KO) rodents have enlarged lamellar bodies (LBs) in their alveolar epithelial type II cells, although the underlying mechanisms remain unclear. Here we performed proteomic analyses on LBs isolated from *Lrrk2* KO mice and found that the LB proteome is substantially different in *Lrrk2* KO mice compared with wild-type mice. In *Lrrk2* KO LBs, several Rab proteins were increased, and subunit proteins of BLOC-1-related complex (BORC) were decreased. The amount of surfactant protein C was significantly decreased in the bronchoalveolar lavage fluid obtained from *Lrrk2* KO mice, suggesting that LB exocytosis is impaired in *Lrrk2* KO mice. We also found that the enlargement of LBs is recapitulated in A549 cells upon KO of *LRRK2* or by treating cells with LRRK2 inhibitors. Using this model, we show that KO of *BORCS6*, a BORC subunit gene, but not other BORC genes, causes LB enlargement. Our findings implicate the LRRK2-BORCS6 pathway in the maintenance of LB morphology.

## Introduction

Parkinson disease (PD) is a neurodegenerative disorder characterized by the selective loss of dopaminergic neurons in the substantia nigra, Lewy body formation in the remaining neurons, and the impairment of motor functions, including bradykinesia, rigidity, resting tremor, and postural instability (Sveinbjornsdottir, 2016). Leucine-rich repeat kinase 2 (LRRK2) has been identified as one of the most common genetic causes of familial PD (Paisán-Ruíz *et al*, 2004; Zimprich *et al*, 2004). Genome-wide association studies identified the association of the *LRRK2* locus with an increased risk of sporadic PD (Satake *et al*, 2009; Simón-Sánchez *et al*, 2009).

LRRK2 consists of 2,527 amino acids, and contains several functional domains, such as a guanosine triphosphate (GTP)-binding domain and a kinase domain (Civiero *et al*, 2012; Mills *et al*, 2012; Vancraenenbroeck *et al*, 2012). It has been suggested that LRRK2 is involved in autophagy, intracellular vesicle trafficking, inflammatory responses, and synaptic transmission (Araki *et al*, 2018). Recently, small Rab GTPases, including Rab3A/B/C/D, Rab5, Rab8A/B, Rab10, Rab12, Rab35, and Rab43 have been identified as physiological substrates of LRRK2 (Steger *et al*, 2016; Ito *et al*, 2016). LRRK2 phosphorylates these proteins at a serine or threonine residue within their switch II domain, thereby regulating the interaction of Rab proteins with their regulatory factors as well as effector proteins. Phosphorylation of these Rab proteins by LRRK2 has been shown to regulate various cellular functions, including the regulation of primary cilia, lipid storage, and the homeostasis of stressed lysosomes (Yu *et al*, 2018; Eguchi *et al*, 2018; Steger *et al*, 2017).

LRRK2 is highly expressed in the brain, kidney, lung, and immune cells (Giasson *et al*, 2006). Although *Lrrk2* knockout (KO) mice did not show any notable changes in the brain, a substantial enlargement of secondary lysosomes in renal proximal tubule cells and lung lamellar bodies (LBs) in alveolar epithelial type 2 (AT2) cells was observed (Herzig *et al*, 2011). Moreover, mice, rats, as well as nonhuman primates administered with selective LRRK2 kinase inhibitors showed a similar enlargement of LBs in AT2 cells (Fuji *et al*, 2015; Harney *et al*, 2020; Andersen *et al*, 2018). These observations led us to hypothesize that LRRK2 plays an important role in regulating LBs. LB enlargement was observed in *Rab38* KO mice, presumably due to a decrease in LB exocytosis (Osanai *et al*, 2010). However, the molecular mechanisms underlying the enlargement of LBs in *Lrrk2* KO mice remains unknown.

Pulmonary surfactant is a mixture of proteins and lipids, and forms a layer on the surface of alveoli to prevent them from collapse during respiration. Lung LBs play an important role in the synthesis, storage, and secretion of pulmonary surfactant (Wadsworth *et al*, 1997). LBs are lysosome-related organelles that exist specifically in lung AT2 cells (Weaver *et al*, 2002). Similar to lysosomes, LBs express lysosomal- associated membrane protein 1 (LAMP1) and CD63, contain soluble degradative enzymes, such as cathepsin C, and have an acidic pH (Hook & Gilmore, 1982). Although the molecular mechanisms of LB exocytosis are not fully understood, lysosomal exocytosis has been relatively well studied. Lysosomal exocytosis requires two sequential steps; i.e., transport to the cell periphery, and fusion with the plasma membrane (Encarnação *et al*, 2016). Recent studies have reported that biogenesis of lysosome- related organelles complex (BLOC) one-related complex (BORC) plays an essential role in the anterograde transport of lysosomes (Pu *et al*, 2015). Given the similarities of LBs to lysosomes, it is possible that LB exocytosis also depends on BORC.

Therefore, in this study we performed proteomic analysis on mouse LBs isolated from wild-type (WT) and *Lrrk2* KO mice. Our results showed that in *Lrrk2* KO mice, several Rab proteins, including Rab3A, Rab3D, and Rab27A were significantly increased, and subunit proteins of BORC, including Borcs6, were significantly decreased. Furthermore, we established a cellular model in A549, a cancer cell line that originated from a human lung, to evaluate LB enlargement. Using these cells, we observed LB enlargement in *BORCS6* KO cells. This effect was rescued by BORCS6 overexpression. Our results hence demonstrated that BORCS6 plays an important role in maintaining the morphology of LBs.

## Results and Discussion

### Lamellar body enlargement in Lrrk2 KO mice

Although several studies have demonstrated that LBs existing in AT2 cells are enlarged in *Lrrk2* KO rodents (Herzig *et al*, 2011), this has not been validated quantitatively. Therefore, we performed electron microscopy on the lung of 2-month-old mice (Figure 1A). In *Lrrk2* KO mice, we found that LBs occupied most of the cytoplasm of AT2 cells, and other organelles were hardly observed. The area occupied by LBs in the electron micrographs was significantly increased in *Lrrk2* KO mice compared with WT mice (Figure 1B). These results indicate that LBs of *Lrrk2* KO mice are significantly larger than those of WT mice. In *Lrrk2* KO mice, the levels of surfactant protein C (Sftpc) in the bronchoalveolar lavage fluid (BALF) measured by the enzyme-linked immunosorbent assay were significantly decreased compared with WT mice (Figure 1C). As Sftpc is secreted by the exocytosis of LBs, this result suggested that LB exocytosis is impaired in *Lrrk2* KO mice.

**Figure 1.**
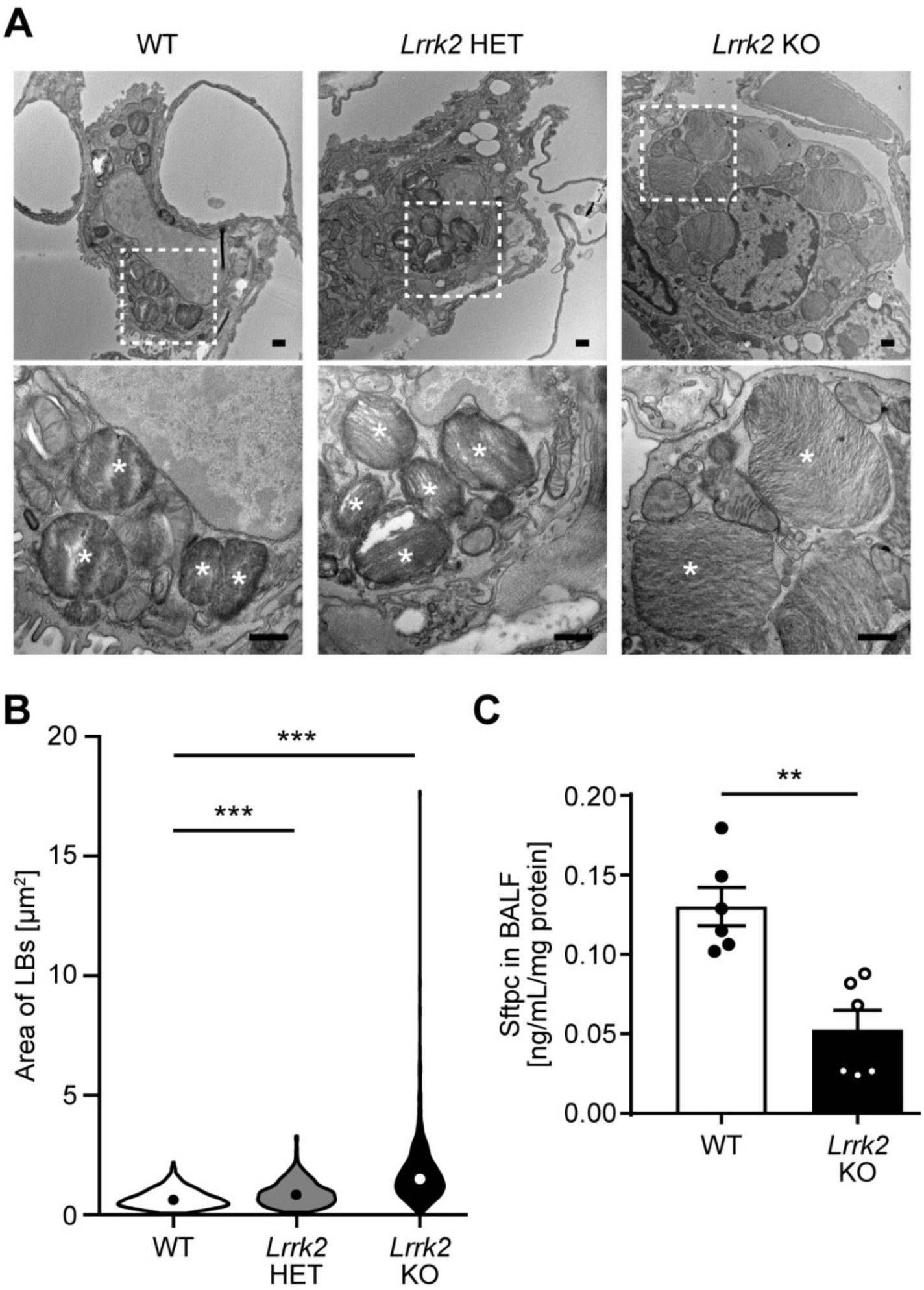
Lrrk2 knockout mice exhibited enlarged lamellar bodies in the lung. A) Representative images of LBs (asterisks in the bottom panels) in WT, *Lrrk2* +/− (HET), and *Lrrk2* KO mouse lungs (male, 2 months old) observed by TEM. Regions marked with white dotted lines in the top panels were magnified in the bottom panels. Scale bars: 500 nm (top/bottom panels). B) The areas of LBs in the TEM images were manually measured on ImageJ and their probability distributions were presented as violin plots. The circles in the plot represent the medians of the values. The total numbers of LBs examined were 382 (WT), 431 (*Lrrk2* HET) and 442 (*Lrrk2* KO). ***p<0.001 (Kolmogorov-Smirnov test). C) The concentrations of Sftpc in BALF collected from WT and *Lrrk2* KO mice were measured by ELISA (n=6, male, 3 months old for both genotypes). The values were normalized by the amount of total proteins in the BALF. The circles in the graph represent individual values. The bars and the error bars in the graph represent the mean values and the standard errors, respectively. **p<0.01 (Student’s t-test).

### Isolation of lamellar bodies from Lrrk2 KO mice

Given that LRRK2 is involved in the regulation of intracellular trafficking, we hypothesized that proteins responsible for the enlargement of LBs might have an altered localization to/from LBs in *Lrrk2* KO mice. To elucidate this hypothesis, we performed liquid chromatography-tandem mass spectrometry (LC-MS/MS) analysis on LBs isolated from mouse lungs. We isolated LBs by sucrose gradient centrifugation (Figure 2A) and confirmed the enrichment of LBs in one of the fractions by immunoblotting (Figure 2B). As a marker of LBs, we used ATP-binding cassette sub-family A member 3 (Abca3), which is an ABC transporter that specifically localizes to LB membranes. We also analyzed the expression of phospholipid-transporting ATPase IA (Atp8a1) as well as lysosome-associated membrane glycoprotein 1 (Lamp1), which are proteins abundantly expressed in the LB membranes (Ridsdale *et al*, 2011). We used Rab5 and receptor-binding cancer antigen expressed on SiSo cells (Rcas1) as markers of early endosomes and the Golgi apparatus, respectively. LB proteins were detected in the 0.4 to 0.5 M sucrose fraction (Figure 2B; lane 8), whereas Rab5 and Rcas1 were not, suggesting that LBs were selectively enriched in this fraction. Notably, LRRK2 was not detected in the LB fraction (Figure 2B).

**Figure 2.**
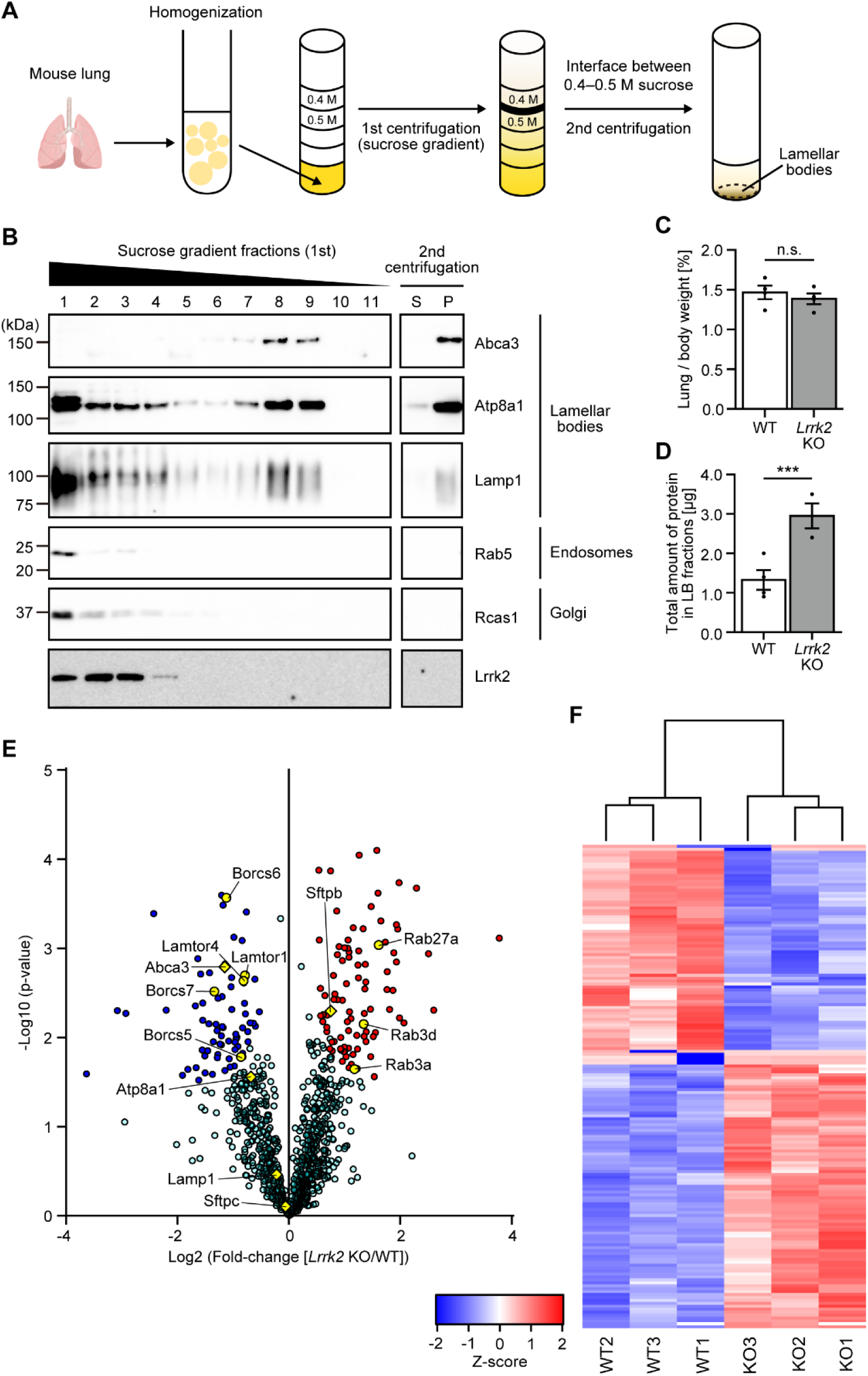
Lamellar body proteome was substantially changed in Lrrk2 KO mice. A) A schematic depiction of LB isolation from a mouse lung by sucrose gradient centrifugation. B) Fractions obtained by the sucrose gradient centrifugation (fractions 1–11 collected from the bottom), as well as the supernatant (S) and pellet (P) fractions after the second centrifugation was analyzed by immunoblotting with indicated antibodies. Abca3, Atp8a1, and Lamp1 are proteins existing in LBs, whereas Rab5 and Rcas1 are markers for endosomes and Golgi apparatus, respectively. C) The percentage of the lung wet weight to the bodyweight of WT and *Lrrk2* KO mice (n=4, male, 3 months old for both genotypes). The dots in the graph represent individual values. The bars and the error bars in the graph represent the mean values and the standard errors, respectively. “n.s.” means “not significant” (Student’s t-test). D) The amount of LB proteins measured on an SDS-PAGE gel stained with SYPRO Ruby. The dots in the graph represent individual values. The bars and the error bars in the graph represent the mean values and the standard errors, respectively. ***p<0.001 (Student’s t-test). E) A volcano plot of the 1,519 proteins quantified in the LC-MS/MS analysis of the LB fractions. Each circle represents a protein. Significantly increased proteins in the *Lrrk2* KO LBs were marked in red, whereas significantly decreased proteins were marked in blue. The proteins picked up in Fig. 3 were marked in yellow circles. The LB proteins analyzed in Fig. 2B were marked in yellow diamonds. F) A heat map of the quantitative values of the LRRK2-regulated proteins. The genotypes (WT1–3 and KO1–3) were shown at the bottom of the heat map. The z-scores were color-coded from -2 (blue) to 2 (red).

### Substantial changes in the lamellar body proteome in Lrrk2 KO mice

Next, we performed LC-MS/MS analysis using the LB fractions isolated from 3- month-old WT and *Lrrk2* KO mice. Although the ratio of wet lung weight to body weight was comparable between the genotypes (Figure 2C), total amount of proteins in the LB fraction was significantly increased in *Lrrk2* KO mice (Figure 2D). The LC-MS/MS analysis identified approximately 1,500 proteins from the LB fraction, and several proteins specifically expressed in LBs, including Sftpb, Sftpc, and Abca3 were shown to be highly enriched, indicating that the LB proteome was successfully acquired (Dataset S1). Label-free quantification demonstrated that 93 proteins were significantly increased by more than 2-fold, and 74 proteins were decreased by less than 0.5-fold in the *Lrrk2* KO mouse LB fraction compared with the corresponding fraction from WT mice (Figure 2E).

### Bioinformatic analyses of the differentially expressed proteins

To further confirm genotype-dependent changes, z-scores were calculated based on the level of each protein quantified in the LC-MS/MS analysis. Clustering analysis based on the z-scores of differentially detected proteins separated the genotypes (Figure 2F). These results suggested that the LB proteome was robustly different between *Lrrk2* KO mice and WT mice.

Gene ontology (GO) enrichment analysis demonstrated that several GO terms, including “small GTPase mediated signal transduction”, were significantly enriched among proteins increased in *Lrrk2* KO LBs (Figure S1A). We also noticed that a large number of Rab GTPases were significantly increased in the *Lrrk2* KO LBs (Dataset S1), some of which have previously been shown to be physiologically phosphorylated by LRRK2 (Steger *et al*, 2017). Proteins decreased in *Lrrk2* KO LBs had GO terms including “negative regulation of peptidase activity”, “anterograde synaptic vesicle transport”, “lysosome localization”, “blood coagulation hemostasis”, “anterograde axonal transport”, and “innate immune response” (Figure S1B). Interestingly, subunit proteins of BORC, including Bloc1s2, Snapin, Borcs5/Loh12cr1, Borcs6/C17orf59, and BORCS7/C10orf32, and subunits of late endosomal/lysosomal adaptor, MAPK and mTOR activator (LAMTOR), including Lamtor1, Lamtor4, and Lamtor5, were significantly decreased in *Lrrk2* KO LBs (Figure 2E, Dataset S1). These results suggested that BORC functions in regulating the size of LBs downstream of LRRK2.

Collectively, based on the results of our proteomic analysis of LBs, we hypothesize that the increase in Rab GTPases and/or the decrease in BORC on LBs are involved in the enlargement of LBs in *Lrrk2* KO mice.

### Altered localization of Rab proteins and BORC subunits on lamellar bodies in Lrrk2 KO mice

To validate the results of the LB proteomics analysis, LB fractions isolated from WT and *Lrrk2* KO mice were subjected to immunoblotting. The expression levels of Rab3A, Rab3D, and Rab27A in *Lrrk2* KO mouse LB fractions were significantly increased, which was consistent with the proteomics results (Figure 3A), whereas no changes were observed in the lung homogenates (Figure 4A). Borcs5 and Borcs7 were significantly decreased in the LB fractions of *Lrrk2* KO mice compared with those of WT mice (Figure 3B), whereas they were unchanged in the lung homogenates (Figure 4B). Unfortunately, we were unable to perform immunoblotting analysis of Borcs6, as all commercially available antibodies for BORCS6 reacted only with human BORCS6, but not with mouse Borcs6. We also confirmed the decrease in Lamtor1 and Lamtor4, which are subunit proteins of the LAMTOR complex, in *Lrrk2* KO LBs compared with WT LBs (Figure 3C), whereas they were unchanged in the lung homogenates (Figure 4C). These results successfully validated the results of our LB proteomics analysis.

**Figure 3.**
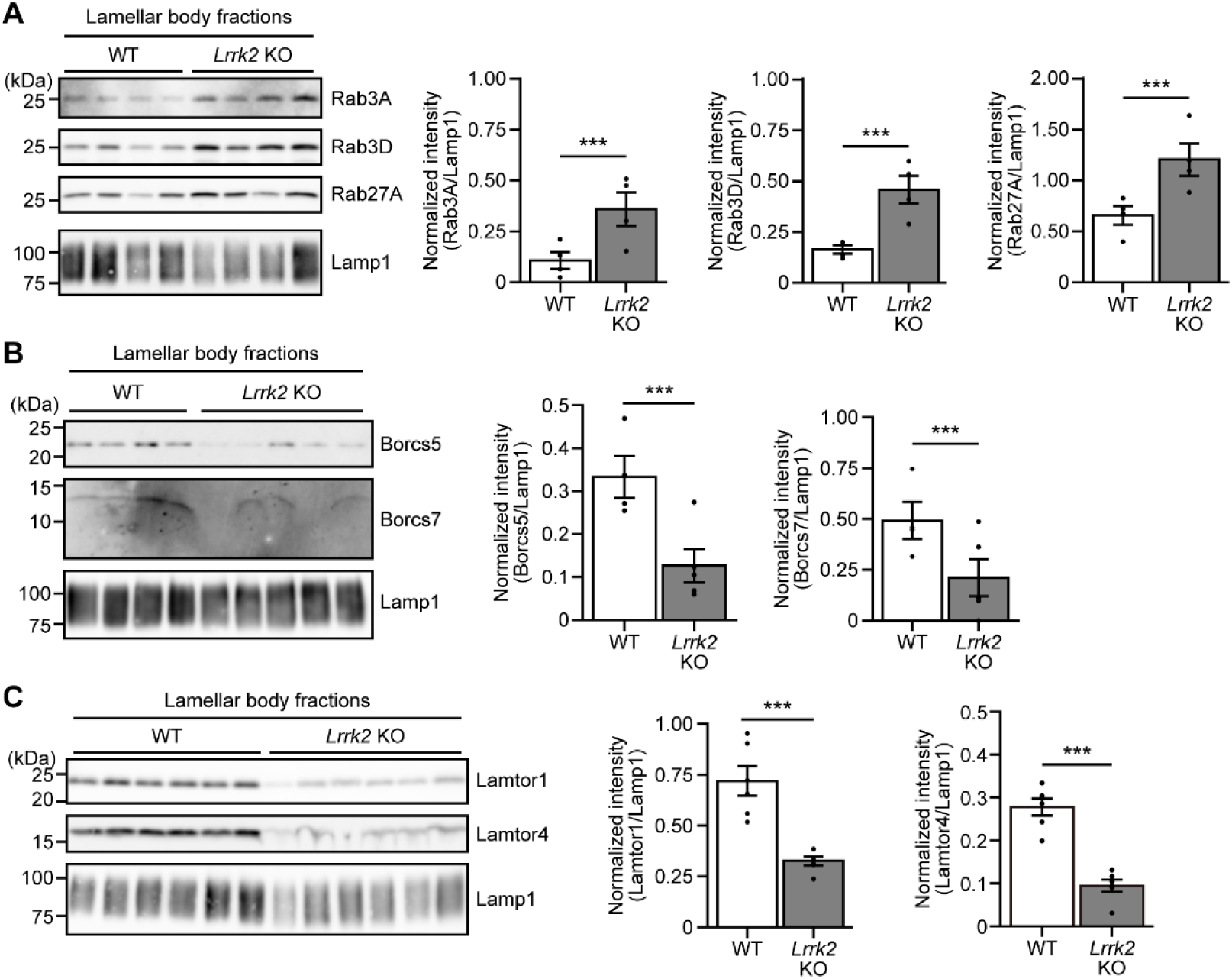
Immunoblot of the regulated proteins in lamellar body fractions. A–C) Expression levels of the indicated proteins in LB fractions were analyzed by immunoblotting (left panel). Lamp1 was used as a loading control. Three-month-old male mice were used for both genotypes. The dots in the graph represent individual values. The bars and the error bars in the graph represent the mean values and the standard errors, respectively. ***p<0.001 (Student’s t-test). The numbers of mice examined were 4 WT and 4 *Lrrk2* KO in (A), 4 WT and 5 *Lrrk2* KO in (B), and 6 WT and 6 *Lrrk2* KO in (C).

**Figure 4.**
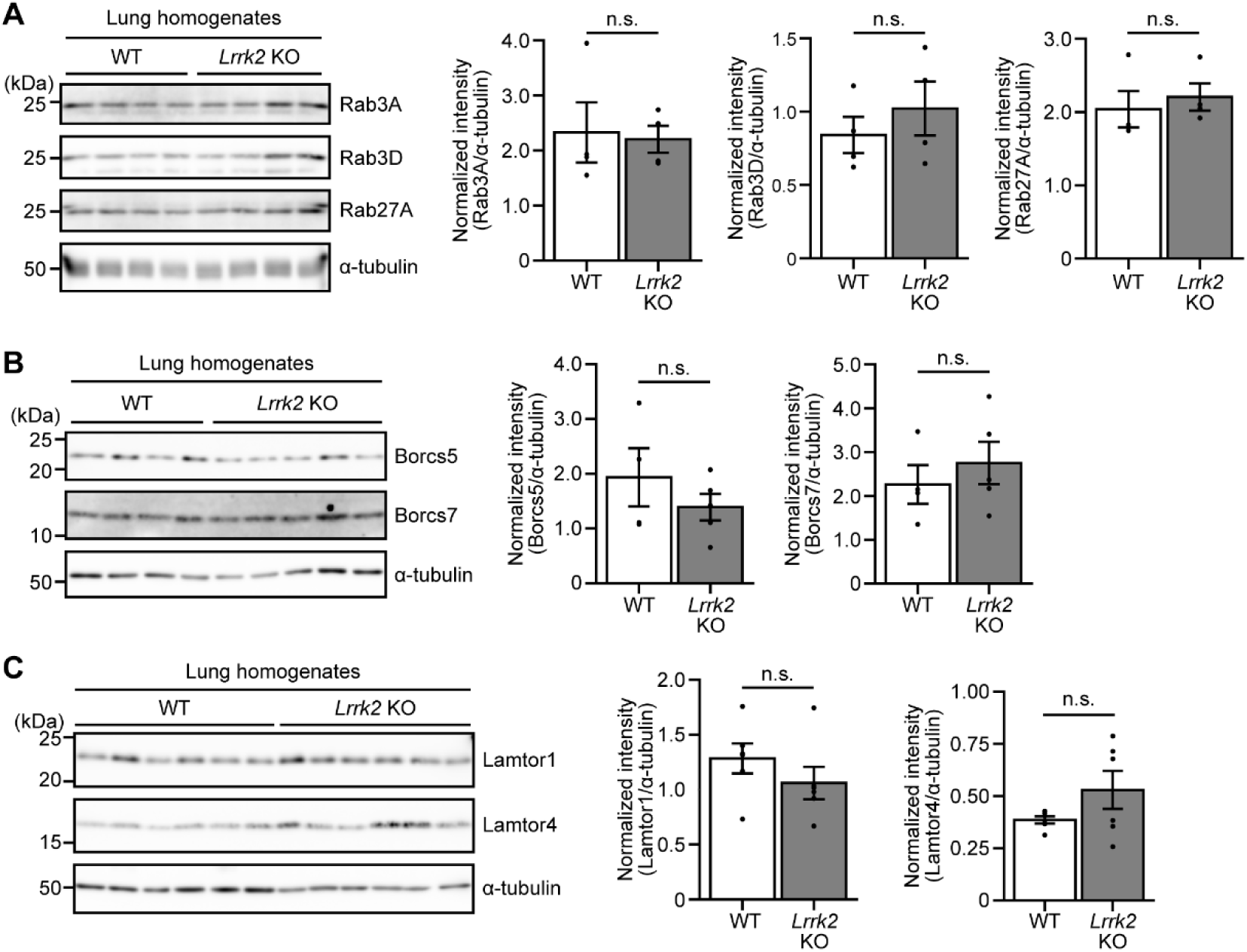
Immunoblot of the regulated proteins in lung homogenates. A–C) Expression levels of the indicated proteins in lung homogenates were analyzed by immunoblotting (left panel). α-tubulin was used as a loading control. Lung homogenates were prepared from the identical sets of mice described in Figure 3. The dots in the graph represent individual values. The bars and the error bars in the graph represent the mean values and the standard errors, respectively. “n.s.” means “not significant” (Student’s t-test). The numbers of mice examined were 4 WT and 4 *Lrrk2* KO in (A), 4 WT and 5 *Lrrk2* KO in (B), and 6 WT and 6 *Lrrk2* KO in (C).

### A cellular model of lamellar body enlargement

To identify the protein(s) responsible for the enlargement of LBs in the absence of LRRK2, we first established a cellular model to analyze LB morphology using A549 cells. As A549 cells were derived from a human lung carcinoma, and harbor multilamellar organelles, these cells have generally been used as a model of AT2 cells (Mason & Williams, 1980). Using the clustered regularly interspaced short palindromic repeats (CRISPR)/Cas9 technology, we generated A549 *LRRK2* KO monoclonal cells (clones #28, #104, and #126). Genomic sequencing confirmed that all clones have indels in the respective genes, resulting in a premature stop codon (Figure S2). Immunoblotting analyses confirmed that these established clones lack the endogenous expression of LRRK2 in contrast to the parental cells (Figure 5A). We also showed that the levels of phosphorylation of Rab10, which is a physiological substrate of LRRK2, were significantly decreased in *LRRK2* KO clones (Figure 5A), indicating the lack of LRRK2 kinase activity in these clones. Electron microscopic observation demonstrated that LBs were enlarged in all *LRRK2* KO A549 clones (Figure 5B). The area occupied by LBs in the *LRRK2* KO cells was significantly larger than that of the parental cells (Figure 5C).

**Figure 5.**
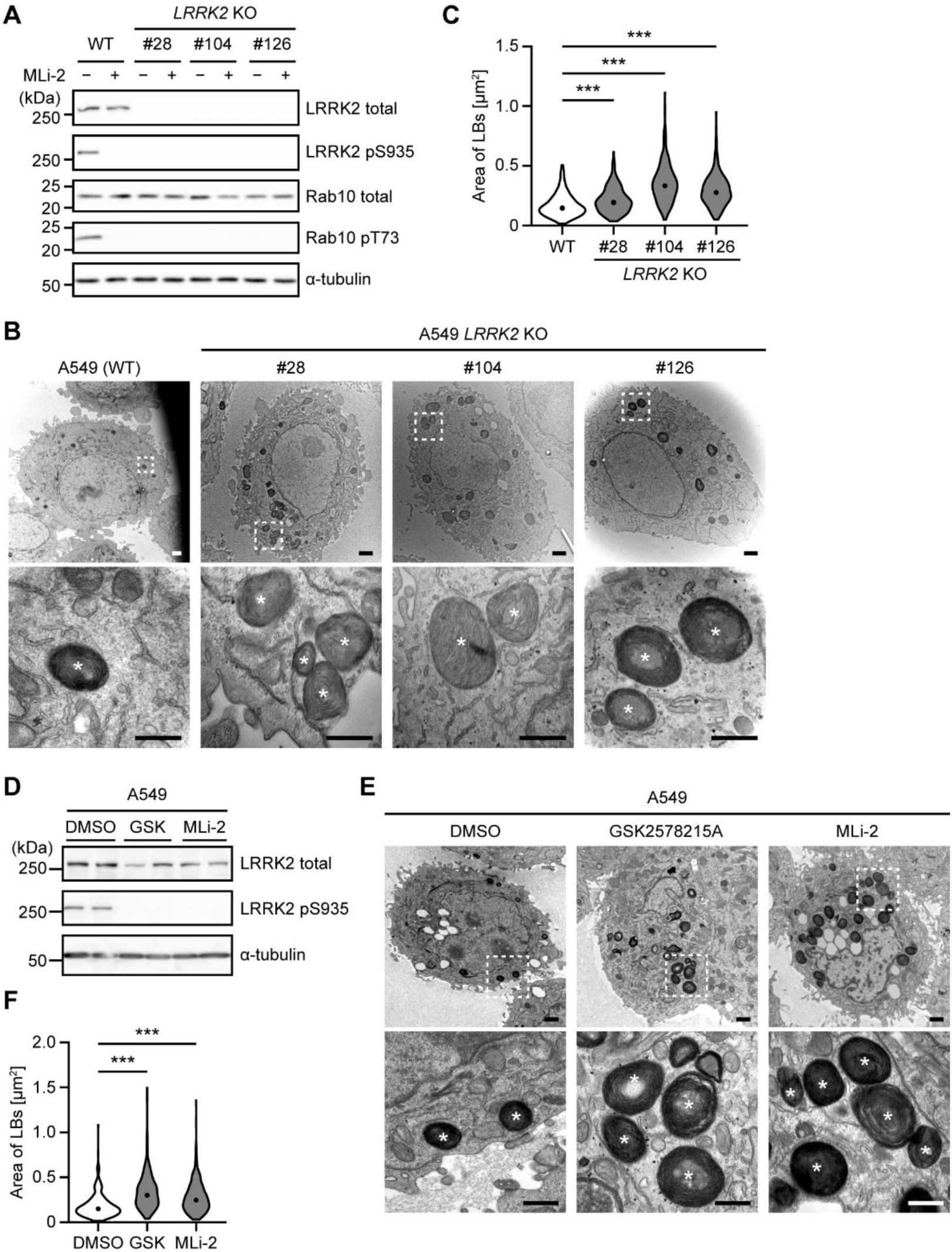
LRRK2 KO caused lamellar body enlargement in A549 cells. A) Expression levels of endogenous total LRRK2, phospho-Ser935 LRRK2, endogenous total Rab10, and phospho-Thr73 Rab10 in A549 parental (WT) as well as in *LRRK2* KO clones (#28, #104, and #126) were examined by immunoblotting. α-tubulin was used as a loading control. Cells were treated with 0.1% DMSO (−) or 10 nM MLi-2 (+) for 24 h prior to lysis. B) Representative images of LBs (asterisks in the bottom panels) in A549 WT as well as *LRRK2* KO cells observed by TEM at low magnification (top panels; scale bars: 1 μm) and high magnification (bottom panels; scale bars: 500 nm). Regions marked with white dotted lines in the top panels were magnified in the bottom panels. C) The areas of LBs in the TEM images were manually measured on ImageJ and their probability distributions were presented as violin plots. The circles in the plot represent the medians of the values. The total numbers of LBs examined were 400 (WT), 398 (#28), 380 (#104), and 401 (#126). ***p<0.001 (Kolmogorov-Smirnov test). D) Expression levels of endogenous total LRRK2 and phospho-Ser935 LRRK2 in A549 WT cells treated with 0.1% DMSO, 1 μM GSK2578215A (GSK), or 10 nM MLi-2 for 1 week were examined by immunoblotting. α-tubulin was used as a loading control. E) Representative images of LBs (asterisks in the bottom panels) in A549 cells treated with DMSO, GSK2578215A, or MLi-2 observed by TEM at low magnification (top panels; scale bars: 1 μm) and high magnification (bottom panels; scale bars: 500 nm). Regions marked with white dotted lines in the top panels were magnified in the bottom panels. F) The areas of LBs in the TEM images were manually measured on ImageJ and their probability distributions were presented as violin plots. The circles in the plot represent the medians of the values. The total numbers of LBs examined were 387 (DMSO), 443 (GSK2578215A), and 470 (MLi-2). ***p<0.001 (Kolmogorov- Smirnov test).

In addition to the genetic model of LB enlargement, we also established a pharmacological model using A549 cells. A549 cells were treated with two selective LRRK2 kinase inhibitors with different chemical structures, namely GSK2578215A and MLi-2, for 1 week. We confirmed that the levels of LRRK2 phosphorylation at Ser935, which is dephosphorylated upon inhibition of LRRK2 by small compounds, were significantly decreased in A549 cells upon treatment with the inhibitors (Figure 5D). The areas occupied by LBs in cells treated with the inhibitors were significantly larger than that of vehicle-treated cells with dimethylsulfoxide (DMSO) (Figure 5E, F). Taken together, we successfully established cellular models to evaluate LB enlargement in A549 cells.

### BORCS6 KO caused lamellar body enlargement in A549 cells

To elucidate whether the decrease in the amount of BORC components on LBs caused the LB enlargement in *Lrrk2* KO mice, we established A549 KO clones lacking either *BORCS5*, *BORCS6*, or *BORCS7*. Genomic sequencing confirmed that all clones have indels in the respective genes, resulting in a premature stop codon (Figure S3). We also confirmed that BORCS5, BORCS6, and BORCS7 were not expressed in the corresponding KO clones by immunoblotting (Figure 6A–C). It has been reported that the KO of BORC components causes the perinuclear accumulation of LAMP1-positive vesicles (Pu *et al*, 2015). Whereas this phenotype was also observed in our *BORCS5* KO and *BORCS7* KO A549 clones, *BORCS6* KO A549 cells did not show the perinuclear accumulation of LAMP1-positive vesicles (Figure 6D). Moreover, the KO of *BORCS5* caused a reduction in the expression level of BORCS7 (Figure 6A) and *vice versa* (Figure 6C), whereas *BORCS6* KO did not affect the expression levels of BORCS5 or BORCS7 (Figure 6B), suggesting that BORCS6 is dispensable for the formation of and function of BORC in A549 cells.

**Figure 6.**
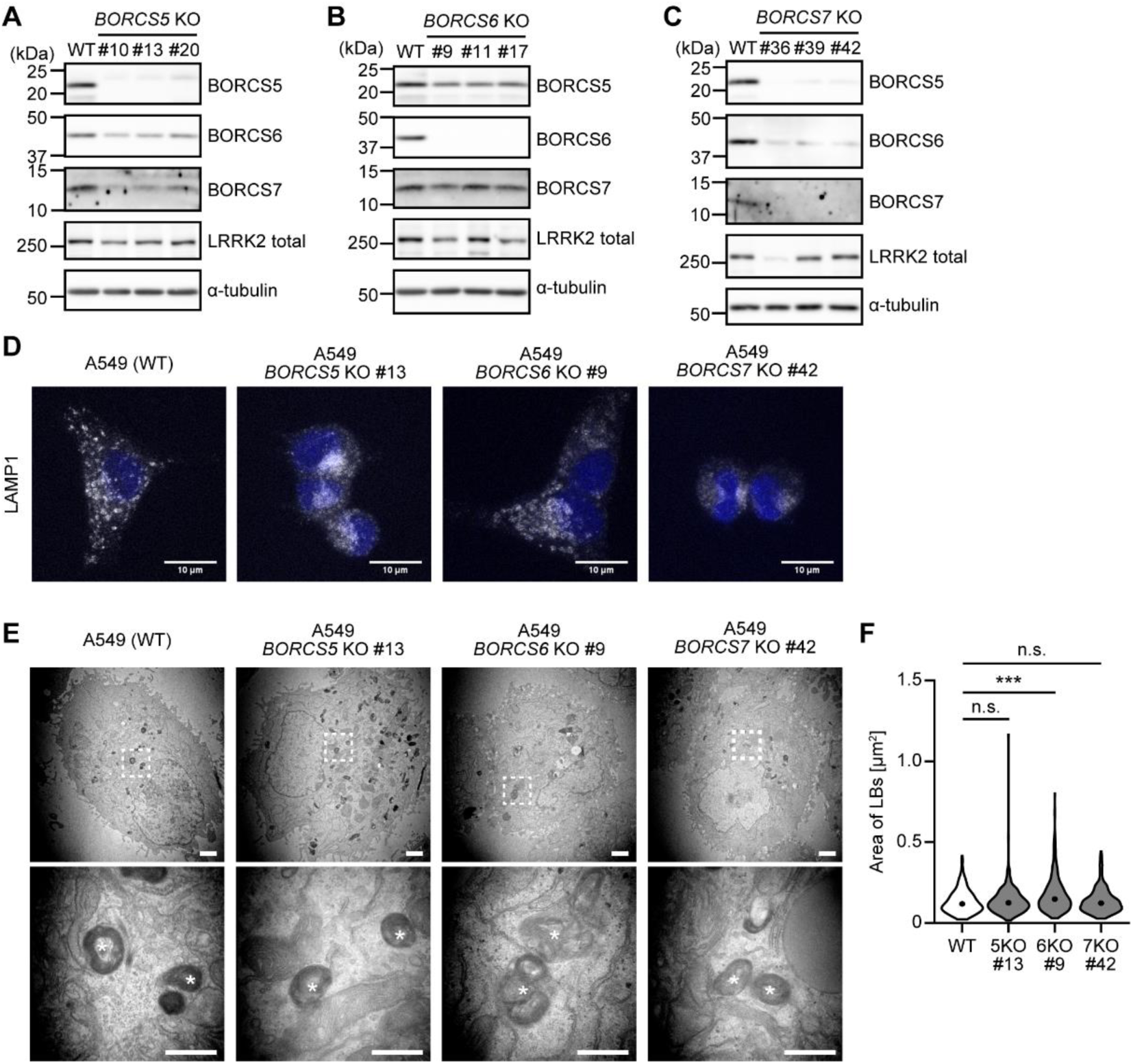
Characterization of A549 cells lacking BORCS5/6/7. A–C) Expression levels of endogenous BORCS5/6/7 and LRRK2 in A549 parental (WT) as well as in (A) *BORCS5* KO clones (#10, #13, #20), (B) *BORCS6* KO clones (#9, #11, #17), or *BORCS7* KO clones (#36, #39, #42) were examined by immunoblotting. α-tubulin was used as a loading control. D) Merged immunocytochemical images of A549 WT, *BORCS5* KO (#13), *BORCS6* KO (#9), *BORCS7* KO (#42) cells stained with an anti-LAMP1 antibody (gray). Nuclei were stained with DAPI (blue). Scale bars: 10 μm. E) Representative images of LBs (asterisks in the bottom panels) in A549 WT, *BORCS5* KO (#13), *BORCS6* KO (#9), *BORCS7* KO (#42) cells observed by TEM at low magnification (top panels; scale bars: 1 μm) and high magnification (bottom panels; scale bars: 500 nm). Regions marked with white dotted lines in the top panels were magnified in the bottom panels. F) The areas of LBs in the TEM images were manually measured on ImageJ and their probability distributions were presented as violin plots. The circles in the plot represent the medians of the values. The total numbers of LBs examined were 426 (WT), 380 (*BORCS5* KO #13), 400 (*BORCS6* KO #9), and 406 (*BORCS7* KO #42). ***p<0.001 (Kolmogorov-Smirnov test).

We next analyzed the size of the LBs in these KO cells by electron microscopy, and found that the *BORCS6* KO clone #9 has enlarged LBs (Figure 6E). Three independent *BORCS6* KO monoclonal clones (*i.e.*, #9, #11, and #17) demonstrated significantly enlarged LBs (Figure 7A, B), but *BORCS5* KO and *BORCS7* KO A549 cells did not show significant changes in the sizes of their LBs (Figure 6E, F). Notably, LBs harboring multiple cores (*i.e.*, multilamellar bodies) were often observed in *BORCS6* KO cells (Figure 6E, Figure 7A). The phosphorylation levels of the physiological substrates of LRRK2, namely, Rab10 and Rab12, were not greatly changed in *BORCS6* KO cells when analyzed by immunoblotting (Figure 7C). Lentiviral overexpression of V5-tagged BORCS6 in A549 *BORCS6* KO cells restored the size of LBs to the level in A549 parental cells (Figure 7D–F), indicating that the on-target deletion of *BORCS6* caused the enlargement of LBs in A549 cells. Taken together, these results suggested that BORCS6 is involved in the enlargement of LBs caused by the absence of LRRK2.

**Figure 7.**
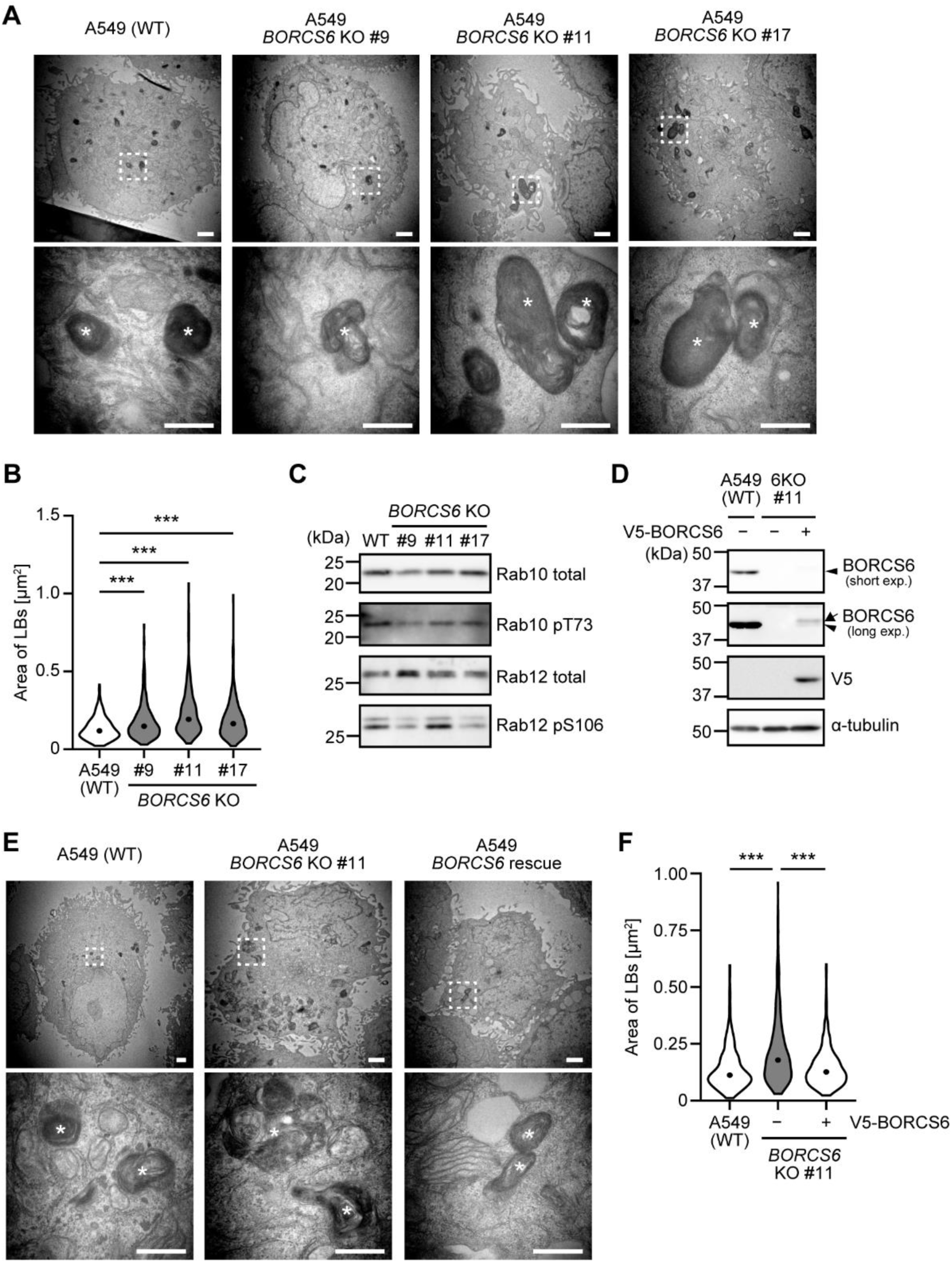
BORCS6 KO caused lamellar body enlargement in A549 cells. A) Representative images of LBs (asterisks in the bottom panels) in A549 WT as well as *BORCS6* KO cells (clones #9, #11, and #17) observed by TEM at low magnification (top panels; scale bars: 1 μm) and high magnification (bottom panels; scale bars: 500 nm). Regions marked with white dotted lines in the top panels were magnified in the bottom panels. B) The areas of LBs in the TEM images were manually measured on ImageJ and their probability distributions were presented as violin plots. The circles in the plot represent the medians of the values. The total numbers of LBs examined were 426 (WT), 400 (#9), 406 (#11), and 417 (#17). ***p<0.001 (Kolmogorov-Smirnov test). C) Expression levels of endogenous total Rab10, phospho-Thr73 Rab10, total Rab12, and phospho-Ser106 Rab12 were examined by immunoblotting. The same sets of samples used in Figure 6B were used. D) Expression levels of BORCS6 in A549 parental (WT) cells, *BORCS6* KO#11 cells, as well as *BORCS6* KO#11 cells stably expressing V5-BORCS6 were examined by immunoblotting. Note that longer exposure was required to detect exogenously expressed V5-BORCS6 (arrow) by an anti-BORCS6 antibody, where the band corresponding to endogenous BORCS6 (arrowhead) in WT cells became saturated (the second panel from the top). α-tubulin was used as a loading control. E) Representative images of LBs (asterisks in the bottom panels) in A549 parental (WT) cells, *BORCS6* KO#11 cells, as well as V5-BORCS6 cells observed by TEM at low magnification (top panels; scale bars: 1 μm) and high magnification (bottom panels; scale bars: 500 nm). Regions marked with white dotted lines in the top panels were magnified in the bottom panels. F) The areas of LBs in the TEM images were manually measured on ImageJ and their probability distributions were presented as violin plots. The circles in the plot represent the medians of the values. The total numbers of LBs examined were 428 (WT), 432 (#11), and 421 (#rescue). ***p<0.001 (Kolmogorov-Smirnov test).

## Discussion

In the present study, we quantitatively analyzed the enlargement of lung LBs in *Lrrk2* KO mice, and systematically identified proteins differentially expressed in the LBs of *Lrrk2* KO mice compared with WT mice, by label-free quantitative mass spectrometry analysis. We found that the LB proteome was substantially different in *Lrrk2* KO mice compared with WT mice, and that several Rab GTPases and BORC subunits had an altered localization in *Lrrk2* KO mice. We then established cellular models of LB enlargement by the KO or inhibition of LRRK2 in A549 cells, and we identified that the loss of BORCS6 causes LB enlargement. These results suggest that BORCS6 is involved in the regulation of the size of lung LBs, and its dissociation from LBs is responsible for the enlargement of LBs in the absence of LRRK2 activity (Figure 8).

**Figure 8.**
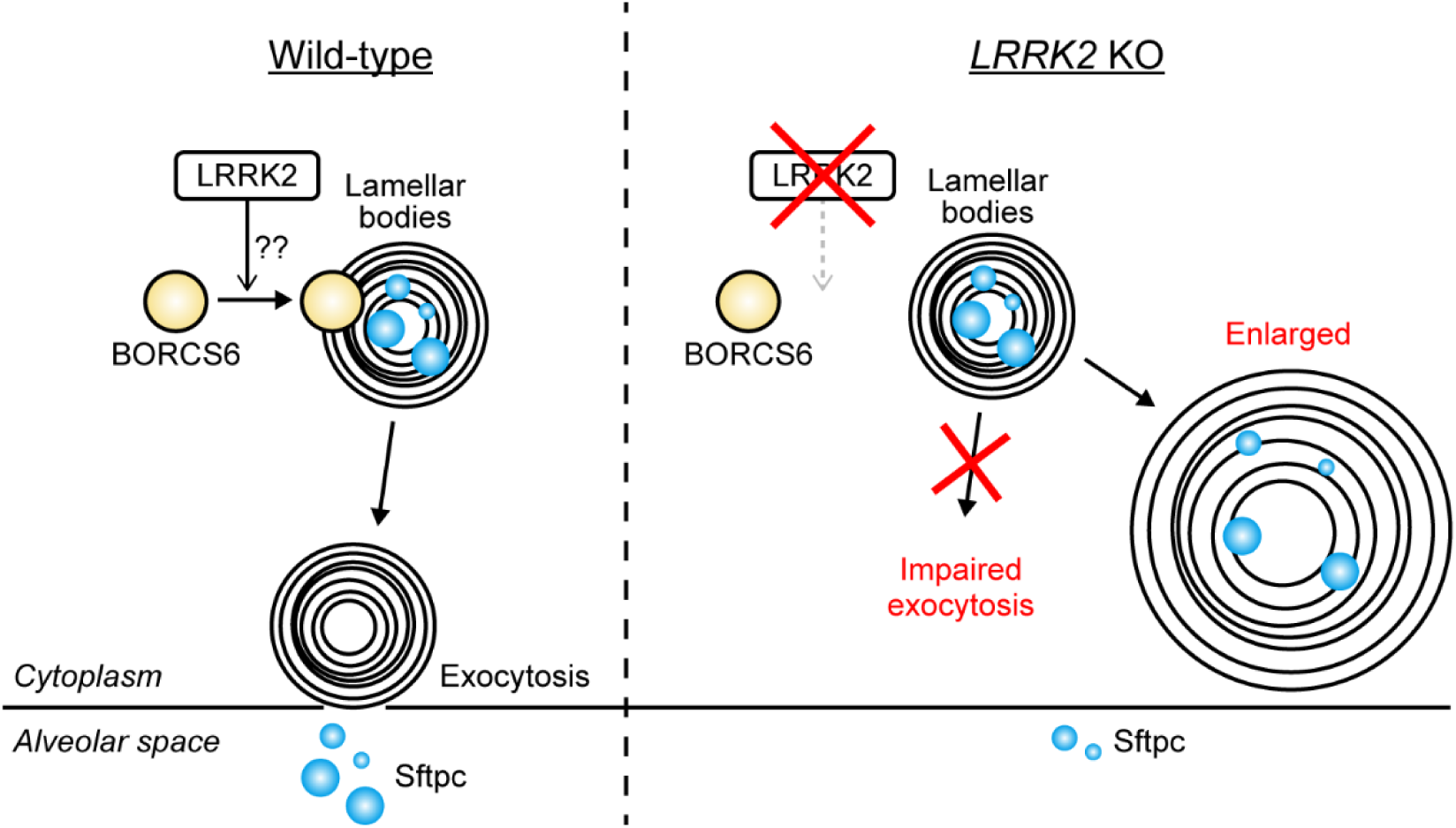
A hypothetical scheme illustrating how LRRK2 deficiency causes lamellar body enlargement through dissociation of BORCS6.

A previous report has shown that AT2 cells primary cultured from *Lrrk2* KO rats are deficient in LB exocytosis (Miklavc *et al*, 2014). In fact, in our study, the levels of Sftpc in BALF were significantly decreased in *Lrrk2* KO mice (Figure 1C), suggesting that the exocytosis of surfactant proteins from LBs are also impaired in *Lrrk2* KO mice. Eguchi and colleagues have shown that in macrophages, LRRK2 promotes lysosomal exocytosis when lysosomes are overloaded with lysosomotropic materials such as chloroquine (Eguchi *et al*, 2018). In this sense, it is reasonable to suppose that in AT2 cells, LRRK2 is also involved in the exocytosis of LBs. It would be interesting to investigate in the future whether the LRRK2-BORCS6 pathway also plays a role in the lysosomal stress response in macrophages.

Among the differentially regulated proteins, several Rab GTPases were significantly increased in the LB fractions of *Lrrk2* KO mice (Figure 2E). These included LRRK2 substrate Rabs, such as Rab3A/D, Rab5A/B/C, Rab8A/B, and Rab10 (Steger *et al*, 2017), as well as non-LRRK2 substrate Rabs, such as Rab1A/B, Rab27A/B, Rab18, Rab6A, Rab31, Rab21, Rab34, Rab33, Rab22A, Rab7A, Rab14, and Rab38. Some of these Rab proteins are involved in the biogenesis of LBs; Rab3D, for example, is localized on LBs at the cell periphery, thereby facilitating their exocytosis (Van Weeren *et al*, 2004). Furthermore, *Rab38* KO animals often show LB enlargement, implicating the involvement of Rab38 in the regulation of LB biogenesis (Osanai *et al*, 2010). However, further investigation is required to unequivocally identify which, if any, Rab(s) play a role in the enlargement of LBs in *Lrrk2* KO mice.

BORC consists of eight proteins, namely, BORCS1 to 8 (Pu *et al*, 2015). BORC exists on lysosomal membranes and promotes the microtubule-dependent centrifugal transport of lysosomes (Guardia *et al*, 2016). Considering that the LB is a lysosome-related organelle and shares some properties with lysosomes, its transport as well as biogenesis may be regulated by BORC. Interestingly, we found that most of the BORC subunit proteins were downregulated in *Lrrk2* KO LBs (Figure 3B), whereas no changes were observed in lung homogenates (Figure 4B). This result prompted us to investigate whether BORC is involved in the enlargement of LBs. We found that KO of the BORC component gene *BORCS6* caused enlargement of LBs in A549 cells, similarly to A549 *LRRK2* KO cells (Figure 6). As the LB phenotype observed upon deletion of *BORCS6* was reproducibly observed in three independent clones, and the phenotype was rescued by the re-expression of BORCS6 (Figure 7), it was clear that the on-target deletion of *BORCS6* caused this phenotype. However, KO of the other two BORC subunits, *BORCS5* and *BORCS7*, did not show similar effects (Figure 6). These results suggest that loss of BORC itself is not involved in the enlargement of LBs observed in the *BORCS6* KO cells. It has been shown that in HeLa cells, the downregulation of BORC components, including BORCS6, causes the accumulation of lysosomes in the perinuclear region (Pu *et al*, 2015; Filipek *et al*, 2017). In contrast, in A549 cells, whereas the KO of *BORCS5* and *BORCS7* caused the perinuclear accumulation of LAMP1-positive lysosomes, the KO of *BORCS6* did not (Figure 6D), indicating that BORCS6 is dispensable for BORC function in A549 cells. These results suggest that BORCS6 may act on its own or in complex with other proteins to maintain the morphology of LBs. It has been shown that BORCS6 (also known as C17orf59 or Lyspersin) associates with LAMTOR on lysosomes, thereby inhibiting the recruitment of mammalian target of rapamycin complex 1 (mTORC1) to lysosomes (Schweitzer *et al*, 2015). As the levels of the subunit proteins of LAMTOR were also decreased in the *Lrrk2* KO LBs (Figure 3C), BORCS6 together with LAMTOR may play a role in the regulation of the size of LBs in AT2 cells.

In summary, we found that BORCS6 is involved in the maintenance of LBs, which we propose to be regulated by LRRK2 (Figure 8). Further studies investigating the molecular mechanism(s) of how LRRK2 regulates the association/dissociation of BORCS6 to/from LBs, as well as how the LRRK2-BORCS6 pathway regulates LB exocytosis in AT2 cells will provide clues towards elucidating the physiological functions of LRRK2.

## Materials and methods

### Animal experiments

All experiments using animals in this study were performed according to the guidelines provided by the Institutional Animal Care Committee of the Graduate School of Pharmaceutical Sciences at the University of Tokyo (protocol no. P29-48). All animals were maintained on a 12 h light/dark cycle with food and water available *ad libitum*. *Lrrk2* KO mice were kindly provided by Professor Jie Shen (Harvard Medical School). PCR genotyping of *Lrrk2* KO mice using genomic DNA extracted from mouse tissues was performed using the following three primers: 5’- GGCTCTGAAGAAGTTGATAGTCAGGCTG-3’, 5’-GAACTTCTGTCTGCAGCCATCATC-3’, and 5’-CTGTACACTGGCAACTCTCATGTAGGAG-3’.

### Quantification of surfactant protein C in BALF

BALF was collected from terminally anesthetized mice by instilling and retracting 1 mL of phosphate-buffered saline (PBS) via a catheter inserted into the trachea. The collected fluid was centrifuged at 500 g for 10 min at 4 °C. The supernatant was used for ELISA and protein assay. The ELISA reaction was performed according to the manufacturer’s instructions. An Sftpc ELISA kit for the mouse was purchased from Aviva Systems Biology (OKEH01170). Total protein concentration in BALF was determined using a Micro Bicinchoninic Acid Protein Assay kit (G-Biosciences; #786-572).

### Isolation of lamellar bodies from mouse lungs

The lungs perfused with PBS were dissected, transferred to the homogenization buffer (1 M sucrose, 10 mM HEPES-NaOH pH 7.5, Complete protease inhibitor cocktail (Sigma-Aldrich), 10 times the volume of the lung wet weight) and homogenized using a Polytron homogenizer (Hitachi) 2 times each for 10 sec on ice. The homogenates were filtered through a 100 µm cell strainer (Falcon), centrifuged at 1,000 g for 10 min at 4 °C to remove cell debris and nuclei, and the supernatants were collected. Sucrose gradient centrifugation was performed using a discontinuous gradient of 0.9 M to 0.2 M sucrose. The post-nuclear supernatants were ultracentrifuged using SW41Ti rotor at 100,000 g for 3 h at 4 °C on Optima L-90K (Beckman Coulter). The fraction between 0.4–0.5 M sucrose was collected, and the sucrose concentration was adjusted to 0.24 M using a refractometer. The samples were then ultracentrifuged one more time at 20,000 g, 15 min at 4 °C to collect the LBs as pellets. For immunoblotting, the pellets containing LBs were solubilized in the SDS–PAGE sample buffer, and the protein concentration of the samples was measured by Micro Bicinchoninic Acid Protein Assay kit (G-Biosciences; #786-572). 2-mercaptoethanol was added to a final concentration of 1% (v/v), and the samples were heated for 15 min at 37 °C.

### Proteomic analysis on isolated lamellar bodies

LBs were isolated from 3-month-old male mice as described above, and the pellets were resuspended in 50 μL per mouse of the lysis buffer (50 mM Tris-HCl pH 8.0, 9 M urea). The suspensions were sonicated 5 times each for 10 sec on ice, and LB lysates were obtained as supernatants following centrifugation. The lysates were snap-frozen in liquid nitrogen and subjected to an LC-MS/MS analysis (Medical ProteoScope, Inc., Japan).

Ten µL of the supernatants were subjected to SDS-PAGE, and the gel was stained with SYPRO Ruby Protein Gel Stain (Thermo Fisher Scientific). Fluorescent images were obtained on an image analyzer LAS-3000 (Fujifilm, Japan). The total protein content of each LB fraction was calculated based on the fluorescence intensity using a known amount of HeLa cell lysate running side-by-side as a standard.

Three hundred μg of the LB lysates were dried and solubilized in a solution containing 8 M urea, 50 mM Tris-HCl, pH 8.0. The cysteine residues were reduced with dithiothreitol at 37 °C for 30 min, followed by alkylation with iodoacetamide. The urea concentration of the sample was adjusted to 2 M using a buffer containing 50 mM Tris-HCl pH 8.0, and mass grade trypsin was added to the samples and incubated at 37 °C for 16 h. Digested peptides were desalted using C18 STAGE tips (Rappsilber *et al*, 2003) and dried under reduced pressure.

The dried peptide samples were dissolved in a solvent (water: acetonitrile: trifluoroacetic acid = 98: 2: 0.1 by volume). The two-thirds of the samples were purified on a nano HPLC capillary column (particle size: 3 µm; inner diameter: 75 µm; length 15 cm) (Nikkyo Technos Co., Ltd., Japan), at a constant flow rate of 350 nL/min, with a gradient 0% to 40% B in 120 min; solvent A: water/formic acid 100:0.1 (v:v); solvent B: water/acetonitrile/formic acid 10:90:0.1 (v:v:v). The MS analysis was performed on a Q Exactive Orbitrap mass spectrometer (Thermo Fisher Scientific) with the top 10 acquisition method: MS resolution 70,000, between 300 and 1500 m/z, followed by MS/MS (resolution 17,500) on the most intense 10 peaks.

Raw MS data were processed using MaxQuant version 1.6.3.3 (Cox & Mann, 2008) with an FDR < 0.01 at the level of proteins and peptides. Searches were performed against the Mouse UniProt FASTA database (downloaded in December 2018). Enzyme specificity was set to trypsin, and the search included cysteine carbamidomethylation as a fixed modification and N-acetylation of protein and oxidation of methionine as variable modifications. Up to 2 missed cleavages were allowed for protease digestion. Quantification was performed by MaxQuant with ‘match between runs’ enabled (matching time window: 0.7 min).

### Bioinformatic analysis on lamellar body proteomes

Bioinformatic analysis for creating the volcano plot shown in Figure 2E was performed on Perseus and data was visualized using Prism (GraphPad Software). The heatmap shown in Figure 2F created and visualized on R, LFQ intensity values calculated using MaxQuant were converted to z-scores by the genefilter package. The heatmap.s function of the gplot package was used to create heatmaps. Gene ontology enrichment analyses were performed using DAVID (https://david.ncifcrf.gov).

### Cell culture

A549 cells (purchased from JCRB cell bank, Japan (JCRB0076)) and Lenti-X 293T cells (Takara Bio, Japan) were cultured in high-glucose Dulbecco’s modified Eagle’s media (DMEM; Fujifilm Wako, Japan; #044-29765) supplemented with 10% (v/v) fetal bovine serum (FBS) (Biosera) and 50 units/mL penicillin and 50 µg/mL streptomycin at 37 °C in a 5% CO_2_ atmosphere. If necessary, cells were treated with GSK2578215A (MedChemExpress), MLi-2 (a kind gift from Professor Dario Alessi (University of Dundee, UK)), or an equal volume of the solvent (dimethyl sulfoxide; DMSO). All cell lines were routinely tested negative for mycoplasma contamination by PCR.

### cDNA cloning and plasmid construction

The cDNA encoding human BORCS6 was cloned from human lung total RNA (Takara Bio, Japan) and inserted into pCR4-TOPO (Invitrogen) by TOPO-TA cloning. The hBORCS6 sequence was amplified by PCR using the following oligonucleotides as primers: 5’-CCTCGGTCTCGATTCTACGGGATCCATGGAGTCGTCT-3’, and 5’- GAGCTCTAGGATATCGAATTCTCGAGTCACTTGCACAGGGCCTCCAACACC-3’ and inserted into the pLVSIN vector (Takara Bio, Japan) by HiFi assembly (New England Biolabs) according to manufacturer’s instructions.

### Lentiviral transduction of A549 cells

A549 cells were plated on 6-well-plate at 5 × 10^5^ cells/well and infected with lentivirus encoding V5-BORCS6. After 24 h incubation, the medium was replaced with a fresh medium. 24 h later, the cells were transferred into a 10 cm dish and cultured with medium containing puromycin at 2 µg/mL. Polyclonal cells obtained after passaging several times were used for rescue experiments shown in Figure 7.

### Generation of CRISPR knockout cells

A549 cells were seeded in 6-well plates at 2 × 10^5^ cells/well and transfected with a set of plasmids targeting a gene (Table S2) using Lipofectamine LTX (Thermo Fisher Scientific) according to the manufacturer’s instructions. At 48 h after transfection, the media were replaced with fresh ones containing puromycin at 2 µg/mL. The media was replaced again at 24 h selection with puromycin. The media were changed to fresh ones not containing puromycin at 48 h selection, and the cells were grown to confluence. For establishing monoclonal cells, the cells after puromycin selection were seeded at a density of 0.4 cells/well into 96-well plates coated with 0.1% (w/v) gelatin (Fujifilm Wako, Japan; #190-15805) and cultured in DMEM containing 30% (v/v) FBS. After reaching approximately 80% confluency, individual clones were transferred to 6-well plates and subjected to immunoblotting. Selected clones lacking the expression of protein-of-interest were sequenced to confirm the knockout: cells were resuspended in QuickExtract (Lucigen), incubated at 65 °C for 15 min, vortexed for 15 sec, and incubated at 98 °C for 10 min. Cell lysates were then centrifuged at 24,400 g for 1 min at 20 °C. Supernatants were collected and PCR was performed using KAPA HiFi HotStart ReadyMix (Roche) to amplify the targeted genomic region. The primers used were listed in Table S3. PCR products were inserted into p3×FLAG-CMV-10 vector (Sigma-Aldrich) by HiFi assembly (New England Biolabs) and transformed into DH5α. Plasmids were isolated from 20 clones using a miniprep kit (Nippon Genetics, Japan) and sequenced to confirm frameshift mutations.

### Immunoblotting

Immunoblotting was performed as described previously (Ito & Tomita, 2017). Antibodies used for immunoblotting were listed in Table S1.

### Immunocytochemical experiments

Cells were cultured on glass coverslips. The cells were washed with DPBS and fixed with 4% (w/v) paraformaldehyde/PBS for 15 min at room temperature. The fixed cells were washed with PBS 3 times and permeabilized in 0.1% (v/v) Triton-X 100/PBS for 30 min at room temperature. The permeabilized cells were blocked in 1% (w/v) bovine serum albumin/PBS for 1 h at room temperature and incubated with the primary antibodies diluted in the blocking buffer overnight at 4°C. After washing with PBS, secondary antibodies labeled with fluorescent dyes were then applied for 1 h at room temperature. The samples were then extensively washed with PBS and mounted using ProLong Diamond (Thermo Fisher Scientific). Images were taken on a confocal microscope (SP5, Leica). Image contrast and brightness were adjusted using ImageJ. Antibodies used for immunoblotting were listed in Table S1.

### Transmission electron microscopy (TEM)

After perfusion of the lung with DPBS, a catheter was inserted into the trachea to wash inside the lung and apply a fixative solution (2% paraformaldehyde, 2.5% glutaraldehyde in phosphate buffer). The lung was removed and incubated in the fixative solution at room temperature with gentle agitation. After 24 h fixation, the lung was chopped into 1-mm^3^ blocks and processed as described below.

A549 cells were seeded in 6-well plates at 6 × 10^5^ cells/well. After 24 h, the cells were washed with DPBS, detached with trypsin/EDTA, and collected into 1.5 mL tubes. Cells were centrifuged at 1,500 g for 5 min at 4 °C and the supernatants were discarded. The cell pellets were washed with DPBS, resuspended in 1 mL of the fixative solution, and incubated for 1 h at room temperature. The cells were then pelleted and washed 3 times with PBS. Five hundred microliters of 4% (w/v) low-melting temperature agarose (Sigma-Aldrich) diluted in double deionized water (DDW) were added to the tubes without collapsing the pellet, and the samples were left to stand at room temperature for 10 min and then on ice for 20 min to harden the agarose. The pellets embedded in agarose were chopped into 1-mm^3^ blocks on ice and immersed in 0.1 M cacodylate-HCl pH 7.4 for 5 min at room temperature. Immersion of the blocks in the cacodylate buffer was repeated 3 times. The blocks were post-fixed with 1% (w/v) OsO_4_ (Nisshin EM, Japan), 1.5% (w/v) potassium ferrocyanide in 0.1 M cacodylate-HCl (pH 7.4) for 1 h on ice. The blocks were then washed with DDW 2 times for 10 min and incubated in 1% (w/v) uranyl acetate (Merck)/70% (v/v) ethanol for 40 min at room temperature in the dark. The blocks were rinsed with 70% ethanol and washed with 80%, 90%, 95%, 99% (once for each), and 100% ethanol for 2 times at room temperature each for 10 min. The blocks were dehydrated twice with QY-1 (Nisshin EM) for 10 min at room temperature, put in the 1:1 mixture of Durcupan (Sigma-Aldrich) and QY-1, and rotated overnight at room temperature. On the next day, the samples were transferred to Durcupan and rotated for 2 h at room temperature. This process was repeated 2 more times. Fresh Durcupan was poured into molds, where the samples were immersed, and the samples were hardened at 60 °C for 48 h. Seventy nanometers-thick ultrathin sections were prepared with an ultramicrotome, picked up onto a mesh (Okenshoji, Japan) covered with Formvar, stained with 4% (w/v) uranyl acetate for 5 min in the dark under a humid condition. After an extensive wash with DDW, the sections were treated with Reynolds solution in the presence of KOH (solid) for 2min. The sections were extensively washed with DDW and imaged under TEM (JEOL-1200EX; JEOL, Japan). Quantification of the area of LBs was performed on ImageJ with the person quantifying kept blind to the sample identity.

### Statistical analysis

Statistical significance of the difference between two samples and among multiple samples was calculated by the Student’s t-test and Tukey-Kramer’s test, respectively, on R. As to the area of LBs, differences of the probability distribution between multiple samples were examined by the Kolmogorov-Smirnov test on R.

## Acknowledgements

We are grateful to Drs. Dario Alessi (University of Dundee, UK) and Jie Shen (Harvard Medical School, USA) for providing MLi-2 and *Lrrk2* knockout mice, respectively. We thank our current and past laboratory members for helpful discussions. Social Cooperation Program of Brain and Neurological Disorders at the Graduate School of Pharmaceutical Sciences, The University of Tokyo is supported by Biogen. This work was supported in part by Biogen; the Grants-in-Aid for Scientific Research (A) [grant number 15H02492/ 19H01015 (to T.T.)]; the Grants-in-Aid for Scientific Research (C) [grant number 17K08265 (to G.I.)]; the Challenging Exploratory Research [grant number 16K15229 (to T.T.)] from the Japan Society for the Promotion of Science; GSK Japan Research Grant 2017 (to G.I.); and the Mitsubishi Foundation (to T.T.).

## Author contributions

MA planned and performed experiments and analyses in all figures, discussed results, and wrote the manuscript. ST was instrumental in experiments involving electron microscopy, and discussed results. GI and TT planned experiments, discussed results, and wrote the manuscript. All authors revised the manuscript.

